# Distinct changes in mature root and growing root tip proteomes underlie physiological responses of bread wheat to salinity stress

**DOI:** 10.1101/2021.05.24.445367

**Authors:** Bhagya M. Dissanayake, Christiana Staudinger, Rana Munns, Nicolas L. Taylor, A. Harvey Millar

## Abstract

The impact of salinity on wheat plants is often studied by analysis of shoot responses, even though the main mechanism of tolerance is shoot Na^+^ exclusion. There is a need to understand the molecular responses of root tissues that directly experience rising NaCl concentrations. We have combined analysis of root growth, ion content and respiration with proteome responses in wheat root tip and mature root tissues under saline conditions. We find significant changes in translation and protein synthesis, energy metabolism and amino acid metabolism in a root tissue specific manner. Translation and protein synthesis related proteins showed significant decreases in abundance only in root tips, as did most of the glycolytic enzymes and selected TCA cycle enzymes and ATP synthase subunits. This selective root tip proteome response indicates protein synthesis capacity and energy production were impaired under salt stress, correlating with the anatomical response of roots and reduced root tip respiration rate. Wheat roots respond directly to soil salinity, therefore shoot responses such as reduction in shoot growth and photosynthetic capacity need to be considered in light of these effects.

## Introduction

Exposure to abiotic stress conditions (low or high temperature, deficient or excessive water, high salinity) cause reductions in growth and development of crops that leads to yield loss worldwide (Wang *et al*, 2003). Among these abiotic stresses, salinity affects approximately 16% of global irrigated land (Flowers & Yeo, 1995). Bread wheat (*Triticum aestivum* L.) contributes nearly 20% of total dietary requirements worldwide and its production is impacted by salinity in many parts of the world, accounting for up to 60% of yield loss which leads to food insecurity in those countries (Dadshani *et al*., 2019). Intensive efforts in breeding salt tolerant wheat varieties have resulted in only limited success in overcoming this problem as salt tolerance is a highly complex process which involves specific morphological, physiological regulatory and metabolic processes (Dadshani *et al*., 2019).

Generally, plants reduce shoot and root growth, photosynthesis and reallocate carbon from growth to maintenance under salinity stress conditions (Jacoby *et al*., 2010). The responses of plants to salt stress occurs in two phases, firstly a rapid response by the growing tissues in the plant to the osmotic stress created by the increase salt concentration in the soil, and then a further and more gradual response as salt concentrations build up in the leaves over time to toxic levels and lead to premature leaf senescence (Munns & Tester, 2008). Increases in the abundance of the enzymes involved in energy metabolism, ROS scavenging enzymes and differential abundance in photosynthesis related proteins (RubisCO subunits, RubisCO activase) has been reported in wheat genotypes under salt stress conditions (Kosova *et al*., 2014).

Recent advances in deployment of ‘omics approaches to crop plants, including proteomic technologies, have provide researchers the opportunity to study the stress responses of plant processes in a comprehensive way that reveals the genes, proteins and metabolites that impact on plant development (Kosova *et al*., 2014; Ahmad *et al*., 2016). However, yield improvement of crops through breeding has mainly focussed on above-ground changes leaving the roots as an under-utilized source for crop improvement. In wheat, proteomic changes in shoot (Jacoby et al., 2013) and root (Wang *et al*., 2008) after salt stress have been reported. A comparison of roots with shoots (Peng *et al*., 2009) observed some common metabolic adaptations. However, these studies were on whole roots and not designed to examine responses of different root types or of different stages of development over time. Root anatomy and physiology both change along the growing root from it tip to more mature regions (Jackson, 2005). The wheat root system responds to saline conditions by an overall decrease in the length of the seminal (axile) roots while the length and number of lateral (branch) roots increase (Rahnama *et al*, 2011). At an anatomical and cellular level, roots respond to salinity by modulating gene expression, protein activities and metabolism resulting in changes in cell wall composition, transport processes, cell size and shape, and root architecture (Byrt *et al*., 2018). Here we have taken into account these physiological responses of the root and have developed untargeted and targeted mass spectrometry approaches to discover, confirm and further explore changes in the root proteome in different wheat roots tissues under salt stress conditions.

## Materials and methods

### Plant materials and growth conditions

Wheat plants (*Triticum aestivum* L. var Scepter) were grown in an unsupported hydroponics system which allows the roots to float freely in the nutrient solution, the system was aerated continuously with air. Upon germination, seeds were transferred to hydroponic solution and grown under controlled environmental conditions described by Che-Othman *et al*., 2020. Salt treatment (NaCl) was commenced after the emergence of the second leaf (5 days post-transplant) (Bado *et al*., 2016). NaCl was added based on the method described by Che-Othman *et al*., 2020. Nutrient solutions were replaced weekly. Harvesting was carried out after 3 days and 6 days of reaching the final NaCl concentration of 150 mM.

### Measurement of biomass root length and ion content

Roots and shoots were separated upon harvest, then he roots were placed in a glass tray and scanned at 600 pixels using a desktop scanner (Epson Expression Scan 1680, Epson America Inc., Long Beach, CA, USA). The obtained images were analysed for root length and average root diameter using WinRHIZO 2008 (model Pro, version 2, Regent Instruments, Canada). Tissue samples were dried in an oven at 70° C for 5 days and the dry weight was recorded. The dried tissue was ground to a fine powder using a Geno grinder and 0.1 g of the resulting powder was added to an acid washed Erlenmeyer flask and 3 mL of 70% (v/v) nitric acid (HNO_3_) was added and allowed the digestion to progress for 20 minutes at 100°C. After the HNO_3_ digestion, further digestion of the material was performed with concentrated (70%) perchloric acid (HClO_4_) at 150° C until the emitted fumes became colourless. The flask was then heated to 170°-180° C, to reduce the volume of the digested solution and was then cooled to room temperature. MilliQ water was then added to dissolve any crystals that had formed. The flasks were then rinsed with MilliQ water three times and added to a vial containing 50 µL of an Yitrium solution (internal standard) and the solution made up to a final volume of 10 mL with the addition of MilliQ water. The Na^+^ and K^+^ content was then quantified using Inductively Coupled Plasma mass spectrometry (ICP-MS) and calibrated using a standard of the ion of interest.

### Measurement of root respiration

The O_2_ consumption of the root tips were measured by an Q2 oxygen sensor (Astec-Global, The Netherlands) in sealed 1.5 mL tubes. The oxygen concentration was measured in 3 minute intervals. The slope of oxygen consumption was calculated after 2 h from the start of the run. The standards 100% N_2_ and normal air were used to calibrate from 0% to 100% atmospheric oxygen, respectively. The oxygen partial pressure was determined to be 20.95% of atmospheric pressure, and the ideal gas law was used to calculate molar oxygen consumption rates (O’Leary *et al*., 2017).

Control and salt treated root tips (1 cm) were harvested after 3 days and 6 days after reaching the final salt concentration (150 mM NaCl). The respiratory rate was measured by immersing the dissected roots in 1 mL of Hoagland solution, pH 6.0, salt treated roots were immersed in the Hoagland solution containing 150 mM NaCl which was exactly same to the hydroponic growth conditions. O_2_ consumption by roots (µmol O_2_ g^-1^ s^-1^) were calculated as the slope between 1 and 3 h after the start of the run. A minimum of seven roots from different plants were analysed in each treatment.

### Root protein extraction digestion, quantification and identification

A 150 mg sample of root tissue was snap frozen and ground with glass beads to a fine powder, then 300 μL of extraction solution (125 mM Tris-HCl pH 7.5, 7% (w/v) sodium dodecyl sulfate (SDS), 0.5% (w/v) PVP40 with Roche protease inhibitor cocktail (Roche) added at 1 tablet per 50 mL) was added. The protein fraction was collected by a chloroform/methanol extraction (Wessel & Flugge, 1984) and the pellet obtained was washed twice with 80% (v/v) acetone. Protein resuspension, quantification and digestion with trypsin was conducted according to the method described by Taylor *et al*., 2014.

Peptide samples were purified using Silica C18 Macrospin columns (The Nest Group) according to instructions provided by the manufacturer and peptides were eluted with 200 µL of 70% (v/v) ACN, 0.1% (v/v) FA solution. The eluate was dried under vacuum and resuspended in 5% (v/v) ACN, 0.1% (v/v) FA to a final concentration of 1 μg/μL. The samples were analysed on a 6550 QToF mass spectrometer (Agilent Technologies, USA) and 6495 triple quadrupole liquid chromatography mass spectrometer (Agilent Technologies, USA).

Peptide mass spectra were submitted to database searching against IWGSC RefSeq 1.0 using MaxQuant program (Version 1.6.1.0). The searching parameters were as follows: 0.5 Da mass tolerance for peptides and 0.01 Da mass tolerance for fragments, carbamidomethyl-cysteine as fixed modification, oxidation of methionine and protein-N-acetylation as variable modifications. The raw mass spectrometry proteomics data have been deposited to the ProteomeXchange Consortium via the PRIDE partner repository with the dataset identifier PXD026000.

### Analysis of un-targeted proteomic profiling

Raw data files generated from Agilent 6550 Q-TOF/LC-MS were analysed by MaxQuant quantitative proteomics software. Statistical analysis was performed using R software (Version 3.6.1) and the Bioconductor package for differentially expressed proteins (DEP 1.1.4 package). This particular R package is specifically developed to analyse mass-spectrometry based proteomics data. The MaxQuant output files were directly used as input files to be analysed by the DEP package. DEP includes features such as, data filtering, normalisation, imputation of missing values and statistical testing. Data filtering was performed according to the following criteria: data must be present for a minimum of two out of three replicates in at least one of the conditions or times tested. After filtering the protein lists, normalisation was performed by variance stabilized transformation (vsn) and data imputation was performed with the knn method which imputes missing values using the k-nearest neighbour approach.

### GO enrichment analysis and KEGG pathway mapping for differentially expressed proteins

In order to identify the metabolic pathways enriched in abundance following salt exposure, hypergeometric-based Gene Ontology enrichment (GO enrichment) was performed with Agrigo software (Du *et al*., 2010) for the differentially expressed proteins identified in both root tips and mature tissues. Specifically, Singular Enrichment Analysis (SEA) was performed by defining IWGSC RefSeq 1.0 as the reference/background proteome, Benjamini-Hochberg (FDR) correction as the method for multiple testing correction and setting 5 as the minimum number of mapping entries per pathway. The adjusted *P*-value (≤0.05) was considered as the threshold to define significantly enriched annotation categories.

In order to observe specific changes in molecular pathways, KEGG (Kyoto Encyclopedia of Genes and Genomes) pathway mapping was performed for the DEP in different tissues at the two different time points. Then the protein sequences were submitted to the BloastKoala tool of KEGG database (https://www.kegg.jp/kegg/kegg1a.html) in order to assign K numbers to the given sequence data by performing a BLAST search. For the BLAST queries “Eukaryotes” were selected as the taxonomy group and used in the scoring scheme for K number assignment. Eukaryotes were also selected as the KEGG database file in order to perform the query. BlastKoala maps the given set of proteins to respective metabolic pathways, these pathways are assigned to main categories; Metabolism, genetic information processing, environmental information processing and cellular processes. The metabolism category includes sub categories such as; carbohydrate metabolism, energy metabolism, lipid metabolism, nucleotide metabolism, amino acid metabolism and metabolism of other amino acids, metabolism of cofactors, biosynthesis of other secondary metabolites and xenobiotics biodegradation and metabolism. Genetic information processing category includes pathways related to transcription, translation and folding, sorting and degradation whereas environmental information processing and cellular processes categories involves signalling pathways and pathways related to cellular processes such as cell growth, cell death and cellular communication respectively.

### Selection and analysis of targeted peptides by Multiple Reaction Monitoring (MRM)

In order to directly determine the abundance of primary metabolism enzymes chosen from the literature, a targeted MRM approach was adopted. Chosen proteins involved in central carbon metabolism or amino acid metabolism in root tissues of wheat were selected from the publicly available functional annotation in the wheat proteome database (Duncan *et al*., 2017). The corresponding protein sequences were imported to Skyline-daily software version 19.0.9.190 (MacLean *et al*., 2010), in the form of a FASTA file, and *in-silico* tryptic peptides were generated by Skyline’s digestion algorithm. Peptide filtering was then applied and included the following conditions: maximum peptide length of 20, each peptide should include at least 8 amino acids, peptides with cysteine and methionine and histidine were discarded in order to avoid the risk of modifications. For peptides with multiple charged fragment ions (doubly or triply charged ions) the ion with the highest intensity for each transition was selected over the other fragment and this was done by comparing the transitions with the reference wheat proteome. For each fragment ion 3 transitions with relatively high intensity were selected and named as rank 1, rank 2 and rank 3 according to their intensities. The resulting peptides were then compared to the entire wheat proteome (International Wheat Genome Sequencing Consortium (IWGSC), 2018) and peptides occurring in proteins annotated as operating in distinct functional bins were removed. As a result, peptides that were selected are unique for a specific functional pathway and can be analysed using triple quadrupole mass spectrometry.

Before conducting MRM analysis on individual experimental samples, a preliminary MRM analysis was performed with a pooled sample which included samples collected at different time points, tissue types and treatments. In order to determine whether the selected peptides could be detected and quantified in different types of tissues. In this dynamic MRM method 211 proteins represented by 1183 transitions were targeted. After obtaining the MRM results for the pooled sample, the final targeted list was refined with the following criteria; at least one peptide per protein and at last three transitions (one quantifier and 2 qualifiers) per peptide. The resulting MRM method contained 892 transitions corresponding to 162 proteins. Raw data files were processed using Skyline (MacLean *et al*., 2010), for each peptide, peak picking was confirmed manually to correct wrongly assigned targets by the Skyline software. Further analysis was performed using peak height of the quantifier ion. Results were normalized by dividing the peak height of each peptide by the average total signal of all targeted peptides in order to correct for differences between the sample runs. Then to give each peptide the same relative importance in protein abundance calculations, the normalised peak height data were scaled by dividing each peptide abundance by the average of that peptide abundance across all the samples.

## Results

### Effect of salinity on wheat biomass, ion content and root respiration

To determine the impact of salt stress on the growth of the plants, the dry weight of shoots and roots were measured 3 and 6 days after reaching the target NaCl concentration of 150 mM in the hydroponic media. Salt stress reduced the dry weight of both roots and shoots to comparable extents (namely by 34% and 28%, respectively) after 6 days of 150 mM NaCl treatment (Fig. **1a,b**). Consequently, there was no significant change in the root:shoot ratio (Supporting information Fig. **S1b**). Chlorophyll concentration of the fully expanded third leaf was reduced by 33% after 6 days of salt treatment (Supporting information Fig. **S1a**). The rate of oxygen consumption by root tips was also significantly decreased during salt stress at both time points compared to the respective controls (by 53% and 36%, on average), indicating the negative impact of salt stress on root respiratory rate (Fig. **1c**). The primary root system showed a significant decrease in total root length and decrease in the average root diameter, consistent with the previous reports and indicating a slowing of root tip growth (Fig. **1d**). (Fig. **1e**, Supporting information Fig. **S1c**). After 6 days of salt stress the Na^+^ concentration in the root was 112 mM (calculated from a FW:DW ratio of 15:1 – see Munns *et al*., 2020) which will require accumulation of organic solutes in order to make up the osmotic adjustment with the external solution. The reduction observed in root growth could be a consequence of this highly energy demanding osmotic adjustment process. Salt treatment reduced the K^+^ level in both roots and shoots (Fig. **1f**, Supporting information Fig. **S1d**), while shoots maintained at least a 10 fold higher K^+^/Na^+^ ratio, even under salt stress conditions, when compared to roots (Supporting information Fig. **S1e,f**).

**Figure 1.**
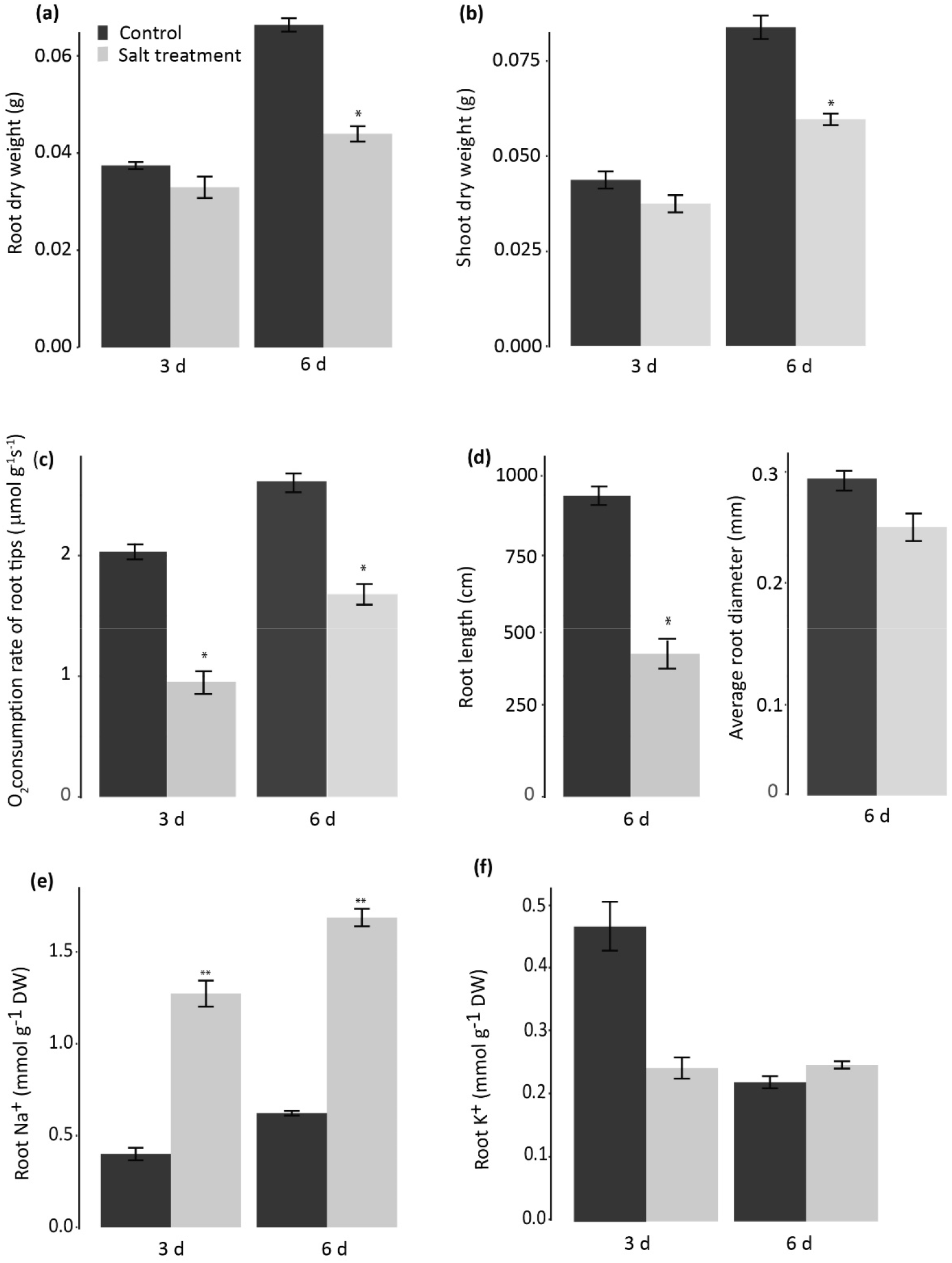
Changes in growth performances, respiratory rates and ion contents of wheat cv. Scepter under 150 mM NaCl treatment for 3 and 6 days. (a) Root dry weight, (b) Shoot dry weight, (c) Rate of oxygen consumption of root tips, (d) Root length after 6 days of salt treatment and Average root diameter after 6 days of salt treatment, (e) Na^+^ content of roots, (f) K^+^ content of roots. Asterisks indicate significant differences between control and salt treatment at each time point separately (Tukey’s Honest Significant Difference; n=4, *, P <0.05)

### Differences between proteomes of mature wheat root and root tips

In order to gain insights into the overall proteome composition of mature roots and how they compared to root tips, an un-targeted proteomic analysis was carried out comparing them. A non-redundant total of 647 and 737 protein groups were detected and quantified in mature roots and root tips, respectively. In both mature roots and root tips the highest number of protein groups were assigned to the functional categories of protein synthesis and degradation in both tissues. Carbon and energy metabolism was the second most abundant functional category, based on number of protein group members, in both tissues (Supporting information Fig. **S2a**). No significant changes were observed in the proportion of identified protein groups assigned to different functional categories in the mature root and root tip proteomes. Among these proteins group sets, 564 protein groups were detected both in mature roots and root tips, while 83 and 173 protein groups were unique to mature roots and root tips, respectively (Supporting information Fig. **S2b**). This showed there is broad similarity in the proteome of the two tissues, but with some quantitative differences in proteome investments.

### Changes of protein profiles in mature roots and root tips during salt treatment

To determine the function of proteins undergoing changes in abundance in different root tissue types during exposure to salinity, un-targeted proteomic analysis was then used to compare salt-treated root tips and mature roots with their respective controls (Table **S1**). In root tips, 50 and 172 protein groups were found to be differentially abundant after 3 days and 6 days of salt stress, respectively, with fold changes greater than 1.5 (*P* ≤ 0.05). In mature roots, 48 and 43 protein groups were differentially abundant after 3 days and 6 days respectively. These protein groups were designated as differentially abundant protein groups (DAPs).

A graph representing the subsets of DAPs found that were significantly increased or decreased in abundance in a particular tissue type and/or at a particular time point is shown in Fig.**2**.

**Figure 2.**
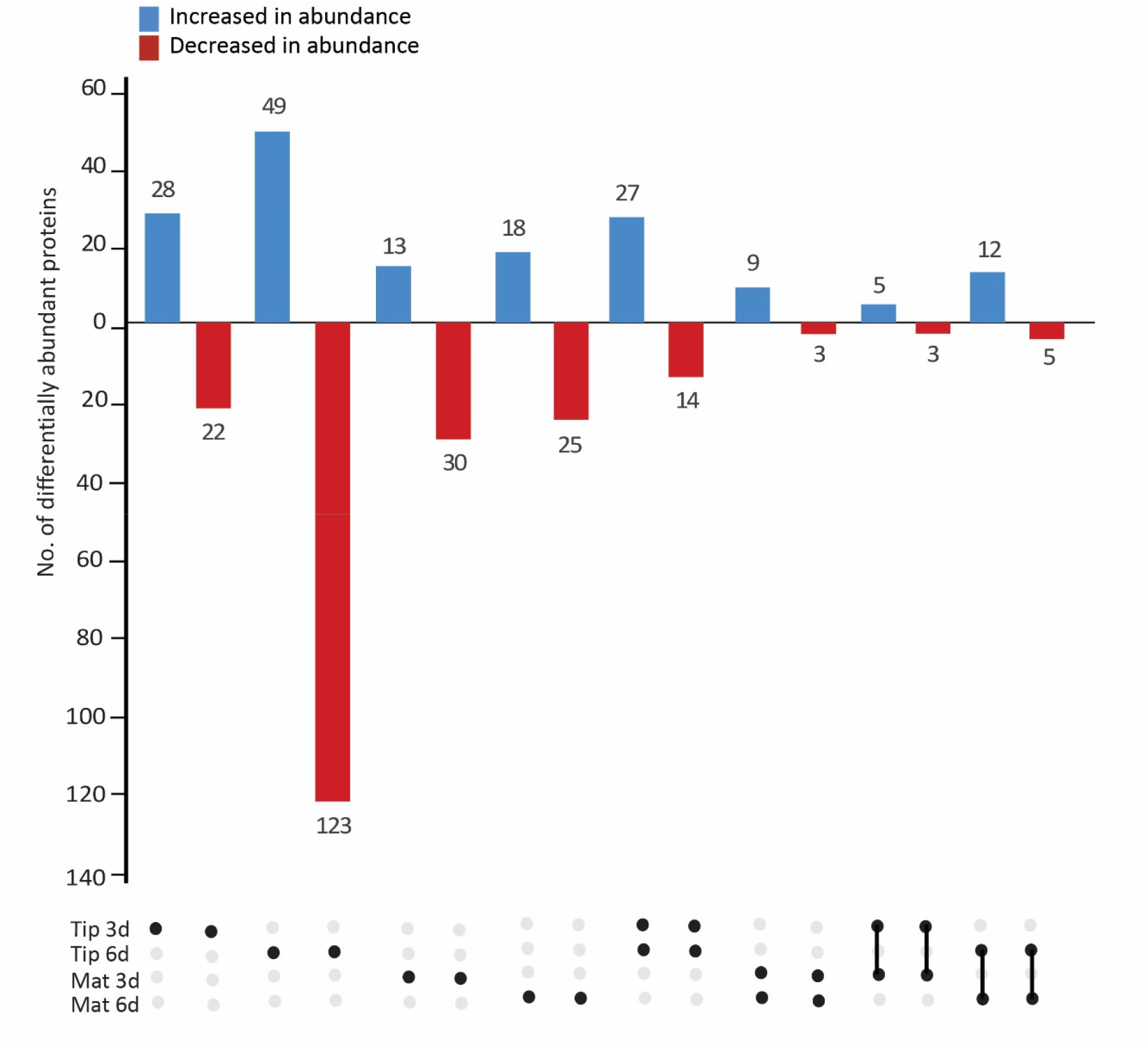
Comparative matrix layout of proteins higher or lower in abundance in different root tissues at different time points. Black dots indicate which sample (Tip 3d, Mat3d, Tip 6d, Mat 6d) are represented by each bar, connected black dots indicates proteins that belongs to different samples of the same root. The number of proteins in the group are indicated on top of each bar. Proteins which increase in abundance are represented in blue colour bars, proteins decreased in abundance are represented in red colour bars.

This illustrates sets of DAPs which were shared by different tissue types at the same time point or shared by the same tissue type at different time points and it provides an insight into the directionality of changes in abundance of shared proteins in response to salinity.

The abundance of proteins found to be differentially abundant in the same root tissues at both time points but potentially differentially abundant in different directions are shown as heat maps in Fig. **3a,b**. In root tips, proteins were identified that are involved in secondary metabolite production, such as alcohol dehydrogenase, phenylalanine ammonia-lyase and O-methyltransferase which all showed increases in abundance under salt stress. Similarly, glutathione S-transferase and peroxidase which are involved in redox reactions and the stress-related protein Endo-1,3(4)-beta-glucanase 1 were found to increase in abundance in root tips, consistent with molecular level changes to cope with salt stress. Proteins that decreased in abundance in root tips were found to be involved in protein folding, development related processes, while proteins involved in protein synthesis and degradation were increased in abundance. In mature roots, peroxidases and proteins involved in secondary metabolism increased in abundance under salt stress. A comparison of different tissue types after 6 days of salt stress (Figure **3c,d**, Fig. **6**) showed that protein synthesis related proteins and proteins involved in cellular organization had an opposing pattern of accumulation in root tips and mature roots with decreases in abundance or increases in abundance, respectively. Interestingly, an aminotransferase involved in the synthesis of alanine was found to be increased in abundance in both tissue types and at both time points (Figure **3c,d**).

**Figure 3.**
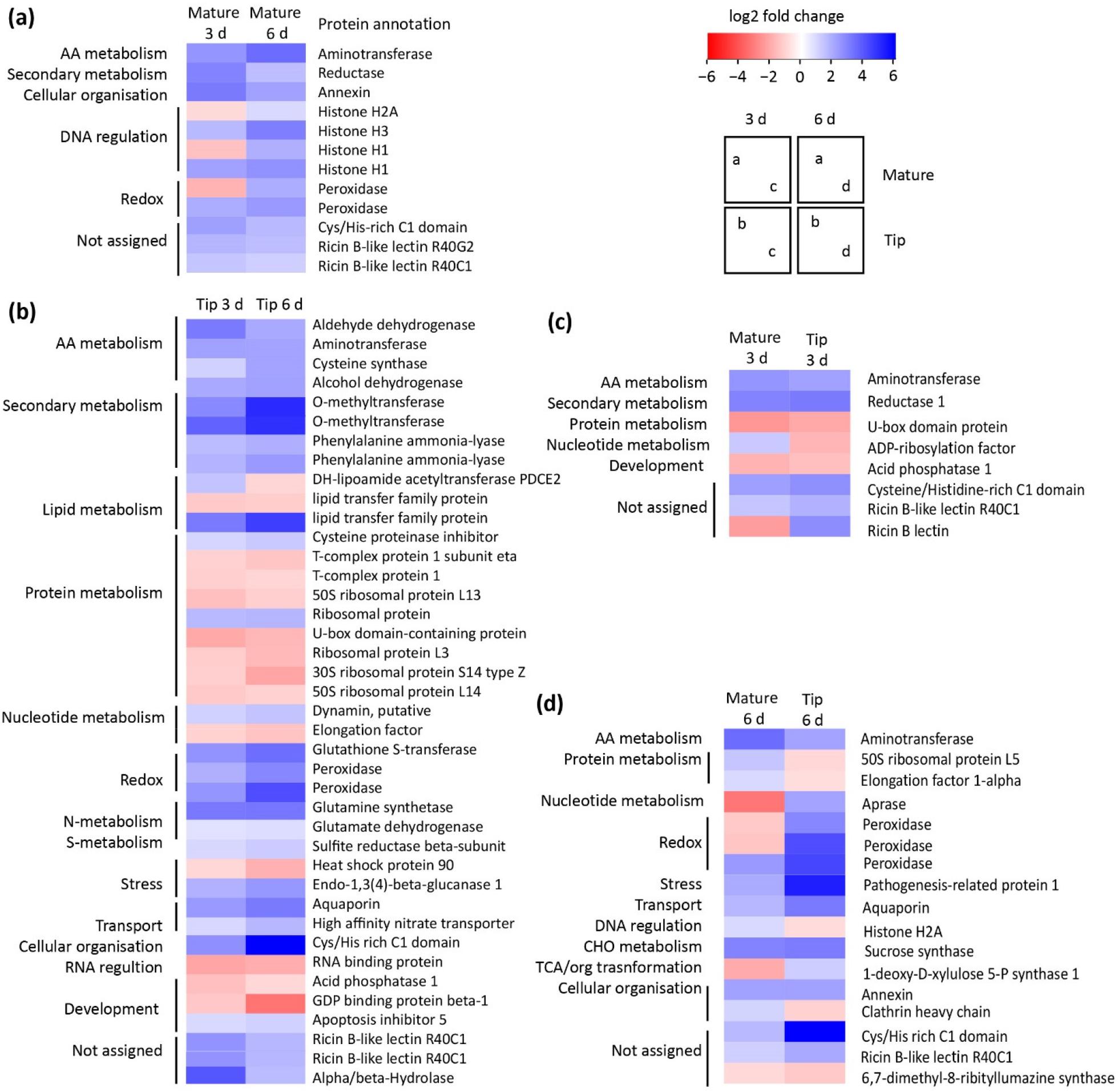
Heat map representing sets of proteins which are differentially abundant (FC>1.5, *P* ≤ 0.05) in mature roots (a) or root tips (b) at both 3 and 6 days of salinity stress, or differentially abundant in both wheat root tissue types after 3 (c) or 6 (d) days of salinity stress.

The majority of the DAPs were found to be differentially abundant in one tissue type at one time point (Fig. **2**), highlighting the spatiotemporal complexity of the response in roots. GO enrichment analysis was performed separately on DAP sets, which revealed that at the 3 day time point, relatively few GO terms were found to be enriched in the DAPs from either mature roots and root tips and GO terms largely failed to provide new insights into the metabolic processes occurring. However, in root tips, DAPs that increase in abundance under salt stress showed a significant enrichment of GO terms “single-organism metabolic processes” and “catalytic activity” while “cellular processes” were significantly enriched in the set of proteins that decreased in abundance under salt stress. Interestingly in mature roots, “metabolic processes” was the only GO term that was significantly enriched in terms of biological processes. In both tissues, GO terms representing cellular components such as “cell” and “intracellular” were found to be enriched (Table **S2**).

In contrast after 6 days of salt stress, DAPs which were found to increase in abundance in root tips showed strong GO term enrichment for “single-organism metabolic process”, “oxidation and reduction process” and “response to oxidative stress”, while “peroxidase activity” (GO:0004601), “oxidoreductase activity” (GO:0016684) and “antioxidant activity” (GO:0016209) were the most enriched molecular functions (Table **S2**). In terms of DAPs which showed decrease in abundance, “translation” (GO:0006412), “nitrogen compound metabolic processes” (GO:0006807), “peptide metabolic processes” (GO:0006518) and “primary metabolic processes” (GO:0044238; eg. glycolytic processes) showed a significant GO term enrichment, whereas “structural molecule activity”, and “structural constituents of ribosome” were enriched GO molecular function terms and “ribosome” was found to be the highest enriched GO term in the cellular component category (Table **S2**).

In mature roots after 6 days of salt stress, “translation”, “peptide biosynthetic process” and “primary metabolic process” showed strong enrichment amongst DAPs, while “structural constituent of ribosome” and “structural molecule activity” were found to be strongly enriched GO terms in biological processes and molecular function categories in DAPs which were found as increased in abundance; “ribosome” also showed the highest enrichment in the cellular components category (Table **S2**). Analysis of DAPs which were decreased in abundance in mature roots revealed enriched GO terms related to response to stress/oxidative stress within biological processes. Notably, peroxidase activity/antioxidant activity showed significant enrichment among molecular function categories; which clearly shows the opposite behaviour of proteins associated with this GO term in mature roots and root tips under the same salt stress condition (Table **S2**). KEGG pathway mapping allowed the observations from GO term enrichment to be placed in a metabolic pathway context. It revealed that the most prominent sets of DAPs from both tissue types occurred in genetic information processing, carbohydrate metabolism, energy metabolism and amino acid metabolism (Fig. **4**). In both mature roots and root tips, a higher number of DAPs were mapped to carbohydrate metabolism and genetic information processing compared to other functional categories (Fig. **4**). However in mature roots, more DAPs in the genetic information category were increased in abundance whereas in root tips a higher number of DAPs were decreased in abundance under salt stress (Figure **4a,b**). Similarly, a considerably higher number of DAPs mapped to energy metabolism, cellular processes and carbohydrate metabolism were decreased in abundance in root tips after 6 days of salt stress (Fig. **4b**).

**Figure 4.**
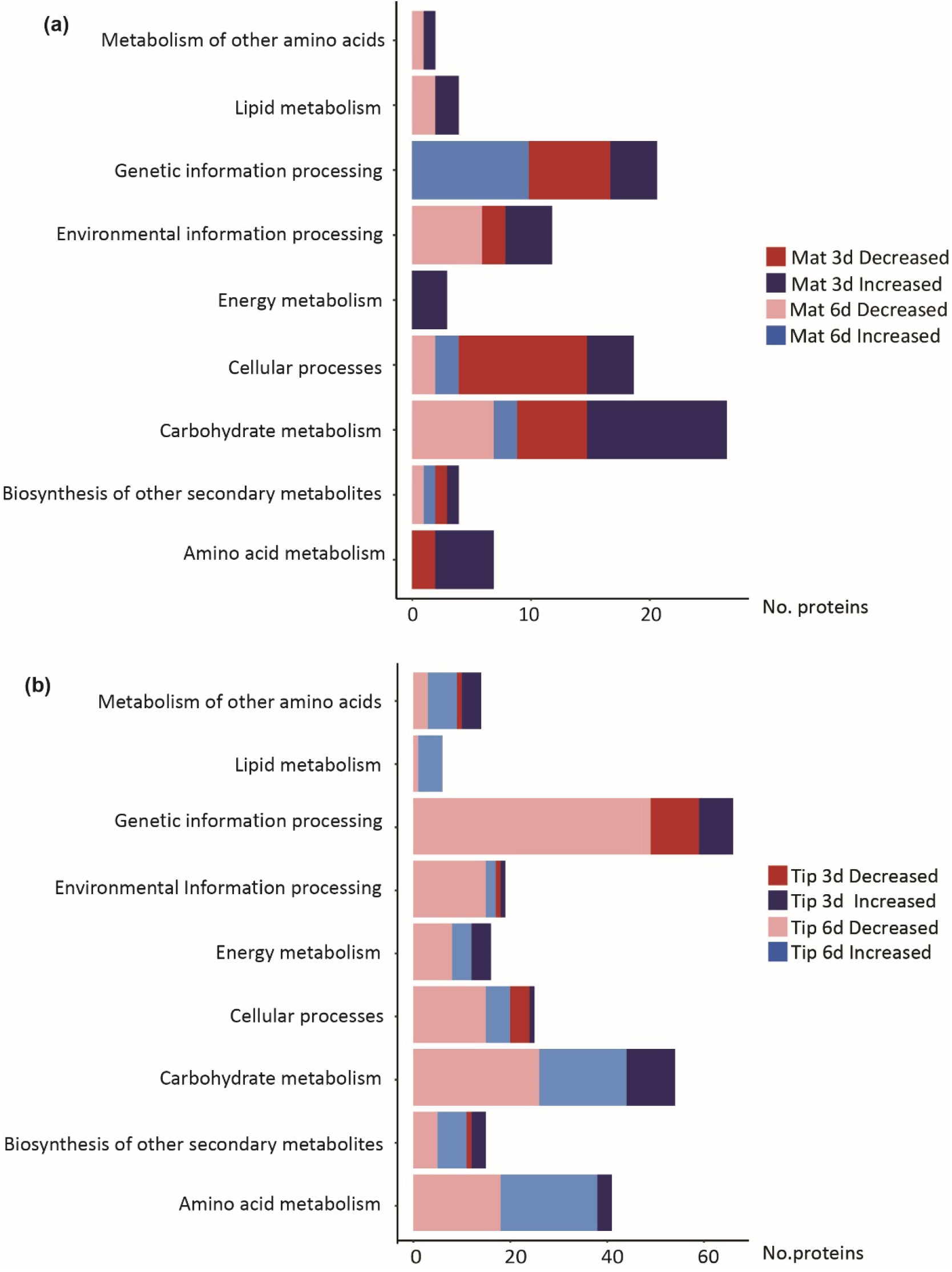
Number of differentially abundance proteins (DAPs) from different functional categories in (a) mature roots, (b) root tips after 3 days and 6 days of salt stress. Colours represents directions of change in abundance of proteins at different time points, dark blue-proteins increased in abundance after 3 days, red-proteins decreased in abundance after 3 days, light blue-proteins increased in abundance after 6 days, pink-proteins decreased in abundance after 6 days. Functional categories were defined by KEGG mapping.

### Targeted analysis of metabolic enzyme abundances in mature wheat roots and root tips

Proteins were then selected to test previous claims of the role of the identified metabolic pathways in salinity whole root studies in other species. A targeted mass spectrometry approach using multiple reaction monitoring (MRM) was used to investigate these responses. Five proteins which were identified and quantified by un-targeted proteome profiling were also quantified using these targeted MRM assays. Two of the protein targets were involved in CHO metabolism (TraesCS7D01G315500: Fructokinase-2, TraesCS7A01G158900.1: Sucrose synthase) and the other three protein targets were involved in glycolysis (TraesCS3B01G087400.1: Triosephosphate isomerase), TCA cycle (TraesCS5A01G505000.2: Aconitate hydratase) and mitochondrial electron transport chain (TaesCS1A01G379000.1: ATP synthase subunit beta). These proteins are indicated in red in Fig. **5**. Similar patterns of abundance were observed for these proteins by both targeted and un-targeted proteomic approaches.

**Figure 5.**
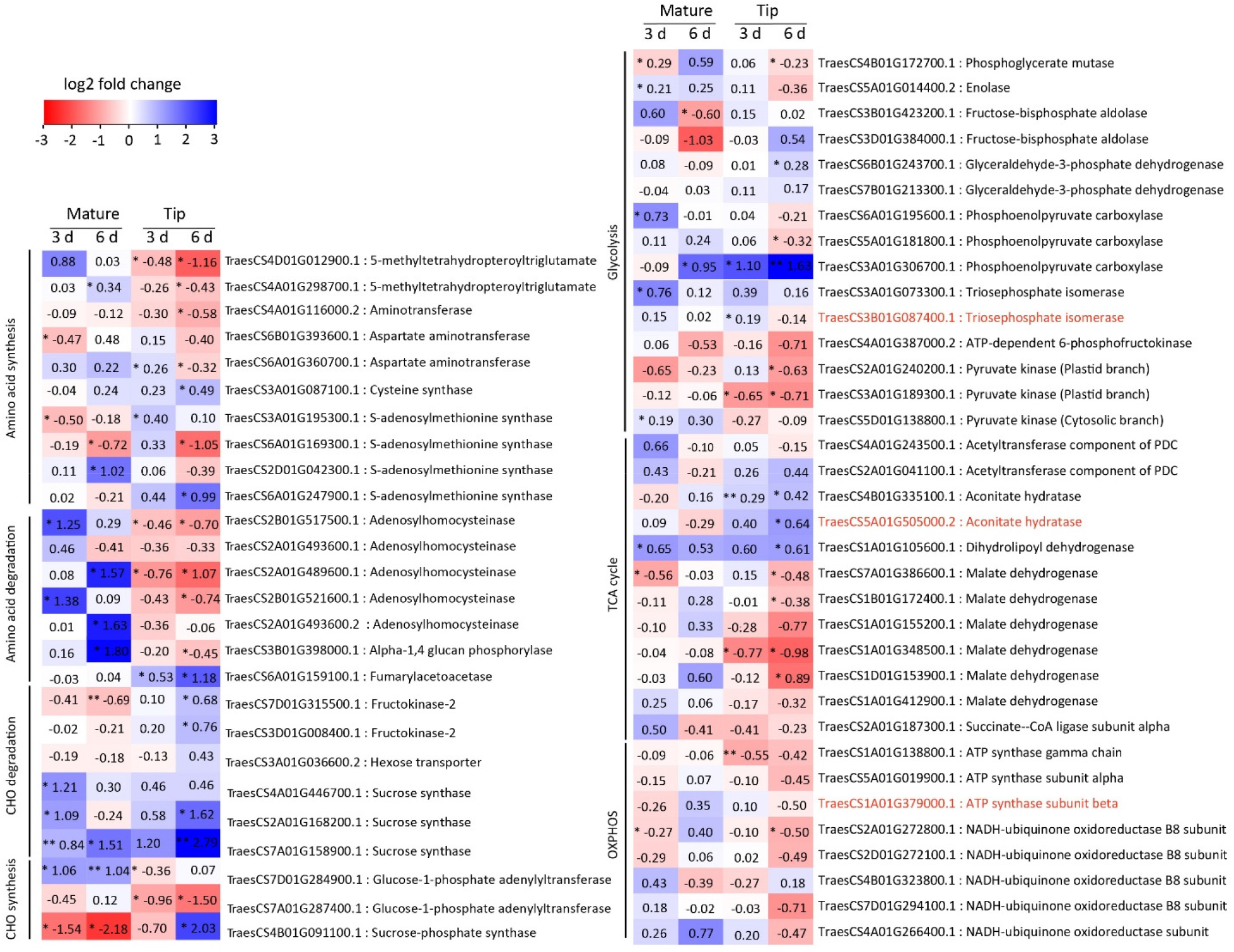
Heat map representing the changes in abundance of wheat root enzymes of selected metabolite pathways during salinity stress. Asterisks indicate significant differences, the red coloured annotations represent proteins quantified by both targeted and un-targeted proteome profiling (t-test for pairwise analysis; * P< 0.05, **, P< 0.01, n=3). The proteins displayed were quantified by analysis of specific peptides using multiple reaction monitoring as shown in Table **S3**.

In the major CHO metabolism category, MRM assays of mature roots showed a lower abundance of sucrose-phosphate synthase which is related to the synthesis of sucrose and an increased in abundance of the sucrose degrading enzyme sucrose synthase at both time points. In root tips after 6 days of salt stress both sucrose synthesis and degradation related proteins showed significantly higher abundance, at the same time enzymes involved in sugar catabolism such as fructokinase-2 were also more abundant (Fig. **5**). The starch synthesis related enzyme, Glucose-1-phosphate adenylyltransferase were also lower in abundance in root tips at both time points under salt stress (Fig.**5**).

Considering the changes of proteins involved in energy metabolism observed in untargeted proteomics (Fig. **3**, Fig. **4**), MRM analysis showed an even larger number of significant changes in abundance for enzymes of glycolysis (Fig. **5**). In mature roots a greater number of significant changes in protein abundance were observed after 3 days of salt stress, but in root tips more significant changes in enzymes in glycolysis and related metabolism were not observed until 6 days of salt stress. However, in root tips after 6 days of salt stress most glycolytic enzymes tested were decreased in abundance except for phosphoenolpyruvate carboxylase, glyceraldehyde 3-phosphate dehydrogenase and triosephosphate isomerase (Fig. **5**). Extending the analysis to pathways that use the glycolytic products, a number of proteins involved in the TCA cycle increased in abundance in root tips after 6 days of salt stress, but significant changes weren’t observed in mature roots (Fig. **5**). Proteins involved in mitochondrial oxidative phosphorylation did not show many significant changes in abundance in both tissues, but in root tips after 6 days of salt stress most of these proteins showed a decrease in abundance (Fig. **5**). The latter is consistent with the slowing of oxygen consumption rate observed in root tips after 6 days (Fig. **1c**).

Glycolytic and TCA cycle intermediates are carbon skeletons for amino acid biosynthesis and products of amino acid degradation. After 3 days of salt stress, targeted quantitation of adenosylhomocysteinase abundance, which is involved in degradation of methionine, showed it was significantly lower and higher in abundance in root tips and mature roots, respectively. Generally, in mature roots, proteins involved in amino acid synthesis such as S-adenosylmethionine synthase and aspartate aminotransferase, were higher in abundance (Fig. **5**, Fig. **6a**). After 6 days of salt stress many significant changes in abundance were observed in root tips compared to mature roots in which most of the proteins identified were involved in amino acid degradation were lower in abundance except fumarylacetoacetase which showed a significant increase in abundance (Fig. **5**), similar to that seen after 3 days of salt stress. Most of the amino acid synthesis related proteins tested showed lower abundance in root tips after 6 days of salt stress (Fig. **5**).

**Figure 6.**
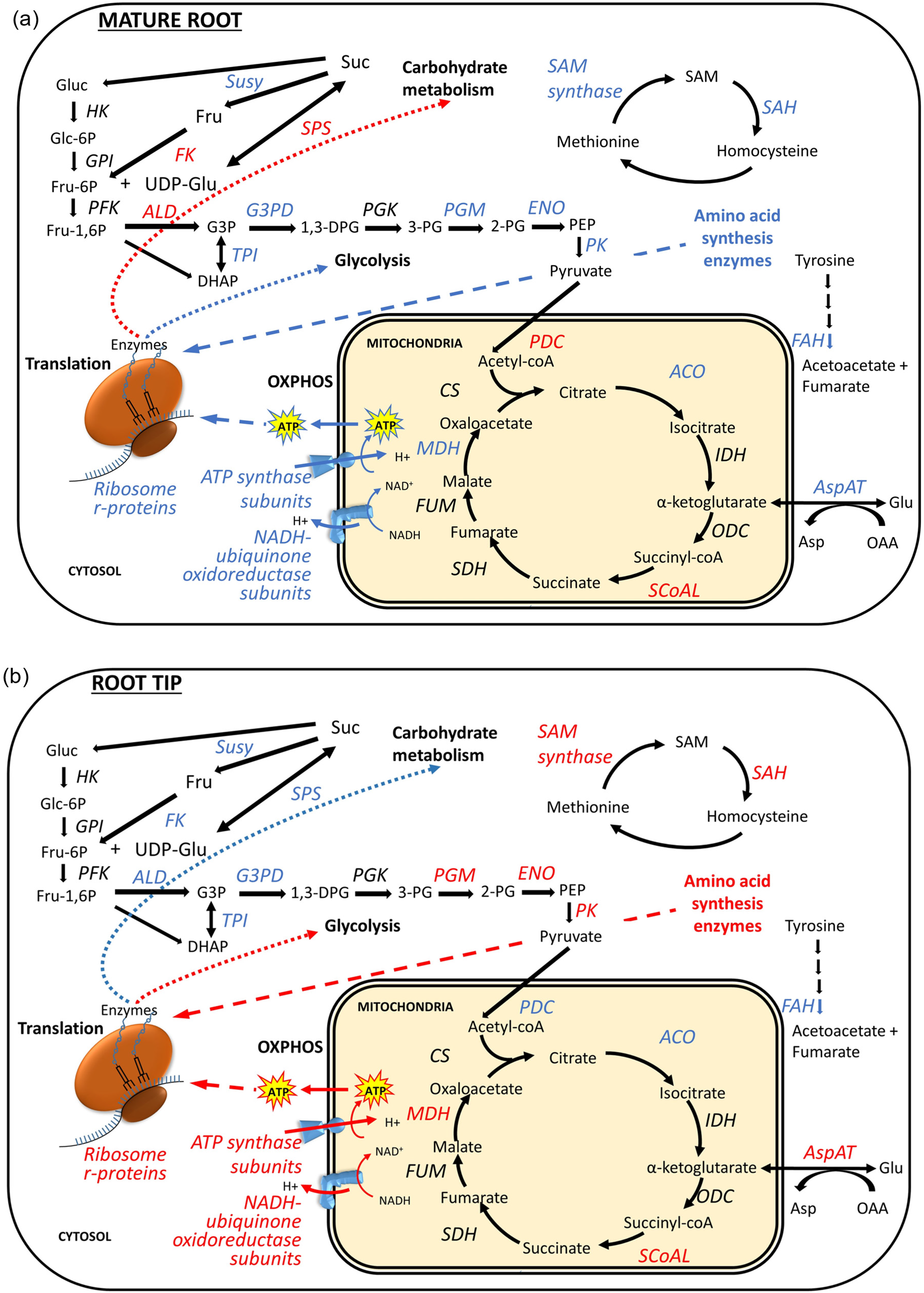
Schematic illustration of proteomic changes observed in (a) mature root and (b) root tips of wheat cv. Scepter after 6 days of salt stress. The influence of cellular provision to ribosomes for protein synthesis is shown by dash arrows (blue increased, red decreased) and the influence of translation on metabolic processes is shown by dotted arrows (blue increased, red decreased). Proteins increased and decreased in abundance are showed by blue colour and red colour respectively, HK-Hexokinase, GPI-Glucose-6-phosphate isomerase, PFK-Phosphofructokinase, ALD-Aldolase, TPI-Triose phosphate isomerase, G3PD-Glyceraldehyde 3-phosphate dehydrogenase, PGK-Phosphoglycerate kinase, PGM-Phosphoglycerate mutase, ENO-Enolase, PK-Pyruvate kinase, PDC-Pyruvate dehydrogenase complex, ACO-Aconitate hydratase, IDH-Isocitrate dehydrogenase, ODC-Oxoglutarate dehydrogenase complex, SCoAL-Succinyl-CoA synthetase, SDH-Succinate dehydrogenase, FUM-Fumarase, MDH-Malate dehydrogenase, CS-Citrate synthase, SAM synthase-S-adenosylmethionine synthase, SAH-S-adenosylhomocysteinase, FAH-Fumarylacetoacetate, ASPAT-Aspartate aminotransferase, Gluc-Glucose, Glc-6P-Glucose-6-phosphate, Fru-6P-Fructose-6-phosphate, Fru-1,6P-Fructose-1,6-bisphosphate, UDP-Glu-UDP glucose, G3p-Glyceraldehyde-3-phosphate, 1,3-DPGA-1,3-bis-phosphoglycerate, 3PG-3-phosphoglycerate, 2PG-2-phosphoglycerate, PEP-Phosphoenol pyruvate, Acetyl-coA-Acetyl coenzyme A, SAM-S-adenosylmethionine, Glu-Glutamate, Asp-Aspartate, OAA-Oxaloacetate, NAD^+^-Nicotinamide adenine dinucleotide, ATP-Adenosine triphosphate

## Discussion

Major commercial Australian bread wheat cultivars like cv. Scepter show a level of tolerance to salinity in the field that enables reasonable harvests even on salt-affected land (Setter *et al*. 2016; Smith *et al*., 2018). Nevertheless, the vegetative growth of cv. Scepter is significantly inhibited by salinity, leading to a reduction of root biomass and root growth (Figure **1a,d**) significant increase in Na^+^ accumulation in both root and shoot tissues, and a decrease of K^+^ concentration. Maintaining a high K^+^ concentration in shoots under salt stress conditions is an important trait of tolerant wheat genotypes (Tester and Davenport, 2003), and the ability to retain K^+^ in root tissues strong correlates with plant salinity tolerance in wheat (Cuin *et al*., 2012). By maintaining a higher K^+^/Na^+^ ratio in shoots compared to the roots, cv. Scepter shows a physiological sensitivity to salt which leads to molecular level changes in different tissues while can be assessed to explore the residual sensitivity or adaptive traits in this cultivar.

Un-targeted proteomic analysis in root tips and mature roots revealed that most of the proteins identified could be assigned to roles involved in protein synthesis and degradation and carbon and energy metabolism. In addition, more proteins were repeatedly detected in root tips compared to mature roots, consistent with the root tip being mainly comprised of young and actively growing cells. Even though the protein sets detected in root tips and mature roots were 68% identical, the subset of these proteins which were found to be differentially abundant under salt stress, varied much more significantly at different time points and in the different tissue types (Fig. **3**). GO enrichment analysis of DAPs in root tips and mature roots revealed that more pathways were enriched in root tips than in mature roots, also when considering the direction of changes in protein abundances of these enriched pathways there was a clear difference between these two tissue types. Translation and protein synthesis related processes were increased in abundance in mature roots, whereas in root tips proteins involved in those processes showed a significant decrease in abundance. Also, enrichment of Gene ontology terms for peptide biosynthetic process and amide biosynthetic process in DAPs decreasing in abundance in root tips further confirms that protein biosynthesis was significantly more affected in root tips than mature roots (Table **S2**). An opposite behaviour amongst proteins related to protein synthesis and processing also occurred (such as elongation factor 1-alpha and 50s ribosomal protein L5 in mature roots and root tips) indicating that salt stress affects the young growing tissue to a greater extent than the mature roots (Fig. **3d**). This can be explained by the reduction of growth as salt exposure restricting the growth of the actively dividing root tissue. Sun *et al*., (2017) showed a significant decrease in abundance of protein isoforms belonging to the 40s, 50s and 60s ribosomal subunits in radish roots treated with 100 mM and 200 mM NaCl which suggests that ribosomal content and potentially protein synthesis capacity of the growing root is impaired by salt stress conditions in that species. However in mature wheat roots studied here, translation and protein biosynthesis subunits increased in abundance under salt stress (Table **S2**) suggesting that older and slower growing root tissues maintain differences in balances between protein synthesis and degradation in order to cope with salt stress conditions.

KEGG pathway mapping showed that most of the enrichment was observed in pathways involved in genetic information processing, carbon and energy metabolism and amino acid metabolism. This confirmed the GO enrichment analysis and showed that the number of proteins related to genetic information processes which showed decrease in abundance was much higher in root tips rather than in mature tissue (Fig. **4**). KEGG pathway mapping further showed that in both root mature and tip tissues the number of DAPs involved in carbohydrate metabolism and amino acid metabolism were larger than the number of DAPs which showed decrease in abundance after 3 days of salt stress. However, as the salt stress continued the number of proteins which showed an increase in abundance were less than the number of proteins which showed a decrease in abundance (Fig. **2**). This showed that the protein abundance changes in root tissues occurred in a time-dependent manner under salt stress conditions. Time-dependent changes were also reported by Witzel *et al*., 2014 in the barley root proteome salt stress response. They showed proteins involved in amino acid metabolism, CHO metabolism and energy metabolism (glyceraldehyde-3-phosphate dehydrogenase, a putative β-1,3-glucanase, a probable plastidic 6-phosphogluconolactonase 5 and asparagine synthetase) markedly change in abundance during the later stages of salt stress.

To schematically illustrate the proteome changes observed in Fig. **3**, Fig. **5**, we have overlaid them to provide an overview of response of root tips and mature roots (Fig. **6**). A greater number of proteins involved in amino acid metabolism were increased than decreased in abundance in both mature roots and root tips. Similarly, many significant changes in amino acid synthesis and degradation related proteins were observed by targeting them directly with MRM assays, further confirming that amino acid metabolism in root tissue was highly impacted by salt stress. In our study, amino acid synthesis related enzymes, aspartate aminotransferase and S-adenosylmethionine synthetase showed significant increase in abundance in mature roots after 6 days of salt stress similar to the responses of radish and barley roots reported by Sun *et al*., (2017) and Witzel *et al*., (2009). However we report wheat root tips showed an opposite pattern of abundance which clearly indicates that different physiological parts of the root system respond differently to the same salt stress condition. Significant GO enrichment of organonitrogen compound biosynthesis and nitrogen containing compound metabolism in DAPs decreasing in abundance in root tips provides more evidence of the significant effect on nitrogen metabolism which occurs under salt stress in roots (Table **S2**). Overall, most of the amino acid synthesis related proteins were found to be significantly decreased in abundance in root tips compared to mature roots which showed an increase in abundance of amino acid synthesis related proteins. This could also indicate that mature roots manage to synthesize amino acids, but root tips favour the degradation of amino acids in order to cope with the impact of salt stress. This degradation is seen to correlate with the lowering of ribosome r-protein subunits and the components of oxidative phosphorylation need to make the ATP required for protein synthesis in root tips (Fig. **6b**).

Carbohydrate and energy metabolism are considered to be important features to fulfil cellular energy demand and to provide essential substrates for other biochemical pathways (Plaxton & Podesta, 2006). A significant increase in abundance in some of the sugar catabolising enzyme sucrose synthase and starch degrading enzymes were observed by targeted MRM assays in mature roots and root tips under salt stress (Fig. **6**) which was also reported by Sun *et al*., (2017). A similar response was also reported by Lakra *et al*., (2019) in rice roots under salt stress. This could indicate that sugar metabolism becomes a more prominent pathway in salt stressed root tissues to generate energy for growth and maintenance.

To maintain energy homeostasis, glycolysis converts energy stored in sugars to ATP and to organic acids that serve as building blocks for amino acids, thereby providing resources to maintain the growth and biosynthesis of plant tissues. By targeted MRM analysis in this study, glycolytic enzymes (except phosphoenolpyruvate carboxylase and glyceraldehyde-3-phosphate dehydrogenase) shown decreases in abundance under salt stress (Fig. **5**, Fig. **6**). Similarly, a notable decrease in many glycolytic enzymes was also reported by Sun *et al*., 2017 in radish roots, indicating a common response of glycolytic capacity being limited under salt stress. Differential expression of glyceraldehyde-3-phosphate dehydrogenase in alfalfa roots and shoots which operate at the end of the TCA cycle to regenerate OAA was reported by (Xiong *et al*., 2017), in which and glyceraldehyde-3-phosphate dehydrogenase showed increased and decreased abundance in shoots and roots respectively. Contrary to this our study showed significant increase in the abundance of glyceraldehyde-3-phosphate dehydrogenase in root tips. Several other studies on transgenic rice roots (Jiang *et al*., 2007) and *Arabidopsis thaliana* roots (Nam *et al*., 2012) have also shown an increase in abundance of glyceraldehyde-3-phosphate dehydrogenase under salt stress. Glyceraldehyde-3-phosphate dehydrogenase is considered to be an indicator of stress tolerance as the increase of it may increase the level of accumulation of soluble sugars in order to provide more energy for plants to cope with salinity (Xiong *et al*., 2017).

The TCA cycle is downstream of glycolysis and is responsible for the oxidation of the substrates to produce most of the aerobic yield of ATP (Sweetlove *et al*., 2010). Four cytosolic malate dehydrogenases (MDH), which operate in the TCA cycle to regenerate OAA, showed a significant lower level of abundance in root tips. MDH was also reported to be decrease abundance in radish roots (Sun *et al*., 2017), similarly decrease in abundance of malate dehydrogenase was also shown in alfalfa roots under salt stress (Xiong *et al*., 2017). However, the prominent increase of abundance in our study of other enzymes at the beginning of the TCA cycle in root tips indicates that this pathway could be more functional under salt stress conditions (Fig. **6b**). One explanation for this is the initiation of an anaplerotic role for the TCA cycle (Che-Othman *et al*., 2017) in which is delivers TCA intermediates to other pathways. Interestingly, the increase of abundance of pyruvate dehydrogenase complex subunits and aconitate hydratase observed in root tips under salinity stress is the opposite of what was reported by Che-Othman *et al*., 2020 in wheat shoot mitochondria, indicating differential regulation of the TCA cycle in root and shoot tissues. Salinity stress may increase the energy requirements of plants in order to operate highly energy consuming processes to ensure ion homeostasis, and allow osmotic adjustment and defence against ROS (Zhu *et al*., 2020). Increased abundance of ATP synthase subunits was reported in salt stressed barley and rice roots (Zhu *et al*., 2020; Yichie *et al*., 2019; Liu *et al*., 2014), however in our study ATP synthase subunits were decreased in abundance in root tips (Fig. **5**, Fig. **6**), which indicates the significant effect of salt stress on energy production in wheat.

Significant increase in abundance of enzymes such as peroxidase in both mature roots and root tips after 3 days and 6 days of salt stress indicates the potential importance of redox homeostasis in roots to cope with salt stress (Fig. **3**). Increased abundance of peroxidase under salinity stress was also reported in roots of many different plant varieties including banana, radish and barley (Ji *et al*., 2019, Sun *et al*., 2017, Witzel *et al*., 2014) indicating it’s a common response of plant roots. Significant increase of abundance of O-methyltransferase, glutathione-S-transferase and phenylalanine ammonia lyase detected by un-targeted proteome profiling analysis in root tips indicates the active production of secondary metabolites to cope with salt stress (Fig. **3**). GO enrichment analysis of DEPs revealed that pathways involved in stress responsiveness, such as “oxidation-reduction process” (GO:0055114), “response to oxidative stress” (GO:0006979), “response to stress” (GO:0006950), “peroxidase activity” (GO:0004601), “oxidoreductase activity” (GO:0016684), “antioxidant activity” (GO:0016209) were significantly enriched in proteins which were found to be increased in abundance in root tips and decreased in abundance in mature roots (Table **S2**). This further confirms a key difference between the two tissue types, highlighting that salt stress responsive pathways are induced in a tissue specific manner.

Overall, the proteomic level changes in abundance suggest that salt stress impacts much more on the actively growing root tissue than the mature part of the root. Therefore, to understand root molecular responses to salinity, interpretation of root tissue growth, metabolism and protein synthesis and degradation also need to be integrated (Fig. **7**). The proteomic changes observed here were also clearly time dependent, and this highlights the importance of integrating the changes in root physiology over time in stress-induced omics studies (Fig. **7**). Also, as roots are the first tissue to experience salt and have relatively rapid metabolic, functional and growth responses to it, then shoot responses to salt need to be considered in light of these changes; not just as direct effects of tissue salt accumulation but also as secondary effects of root responses.

**Figure 7.**
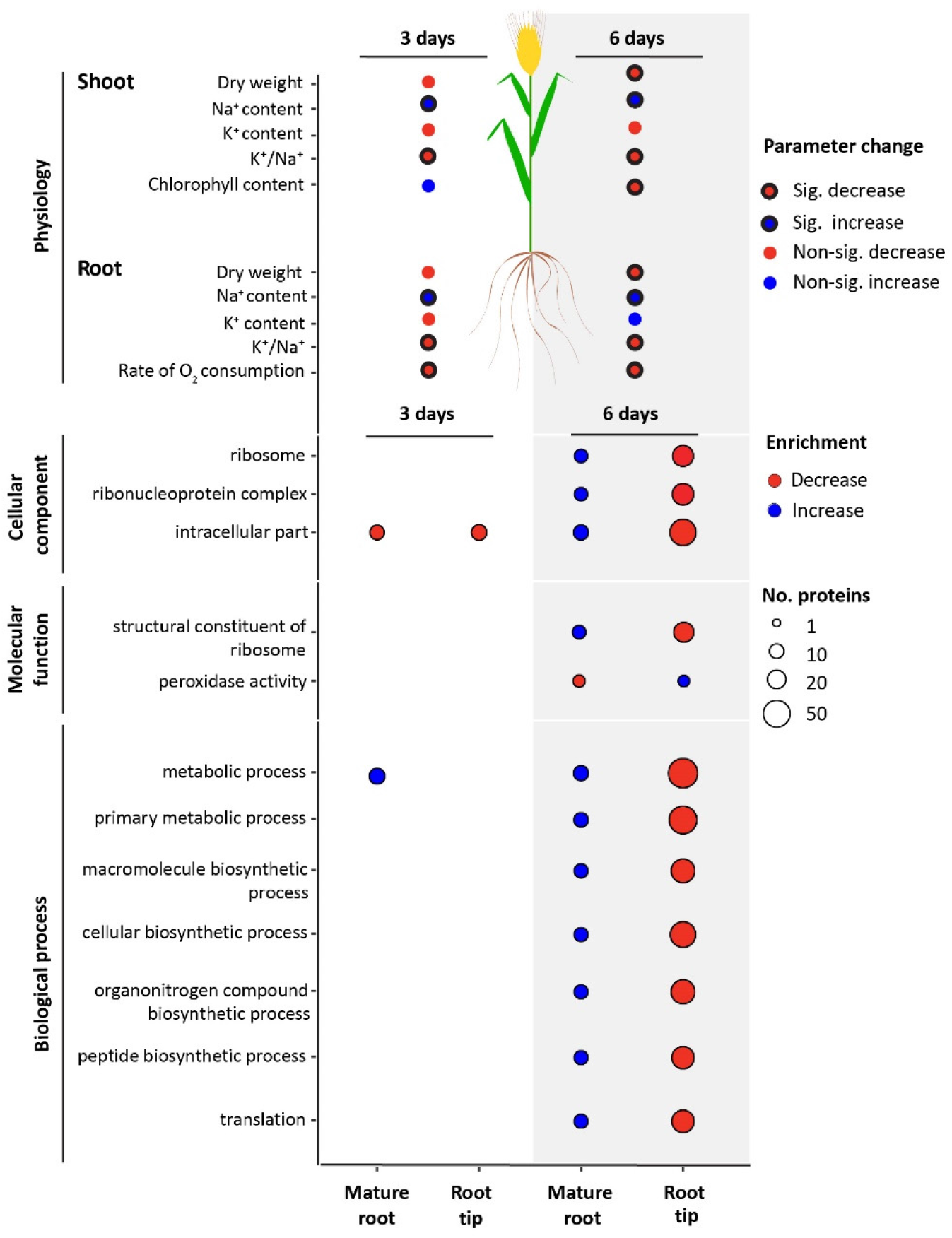
Integrative plot representing physiological and proteomic changes of different root tissues and whole roots and shoots of bread wheat after 3 days and 6 days of treatment with 150 mM NaCl. In the upper panels, red and blue bubbles represent the changes of physiological parameters showing decrease and increase respectively by red and blue colours, the boldness of bubble indicates significance. In the lower panels, bubble plots show the most significantly enriched GO pathways in mature roots and root tips, size of each bubble represents the number of proteins involved in each pathway, colours of the bubbles represent the changes in abundance of proteins involved in each pathway, red - decreased in abundance, blue - increased in abundance.

## Author Contributions

BMD planned and designed research, performed the experiments, undertook data analysis and interpretation, and wrote the manuscript; CS, AHM, RM and NLT participated in planning and designing the research, provided advice on the data analysis and interpretation and contributed to writing the manuscript.

## Acknowledgements

BMD is the recipient of a Scholarship funded by the Grains Research and Development Corporation. This work was supported through funding by the Australian Research Council (ARC) (CE140100008) to AHM. We would like to thank Ms Ricarda Fenske (The University of Western Australia) for supporting the mass spectrometry.

## Supporting Information

**Table S1**. Protein identification and abundances from mature root and root tips after 3 and 6 days of salinity treatment and matched controls.

**Table S2**. Gene Ontology (GO) analysis of differentially expressed proteins (DEPs) by biological process (BP), molecular function (MF) and cellular component (CC) categories in wheat mature roots and root tips in response to salinity treatments.

**Table S3**. Quantification of the abundance of peptides for specific enzymes in different metabolic pathways in wheat mature roots and root tips in response to salinity treatments through multiple reaction monitoring.

**Figure S1.**
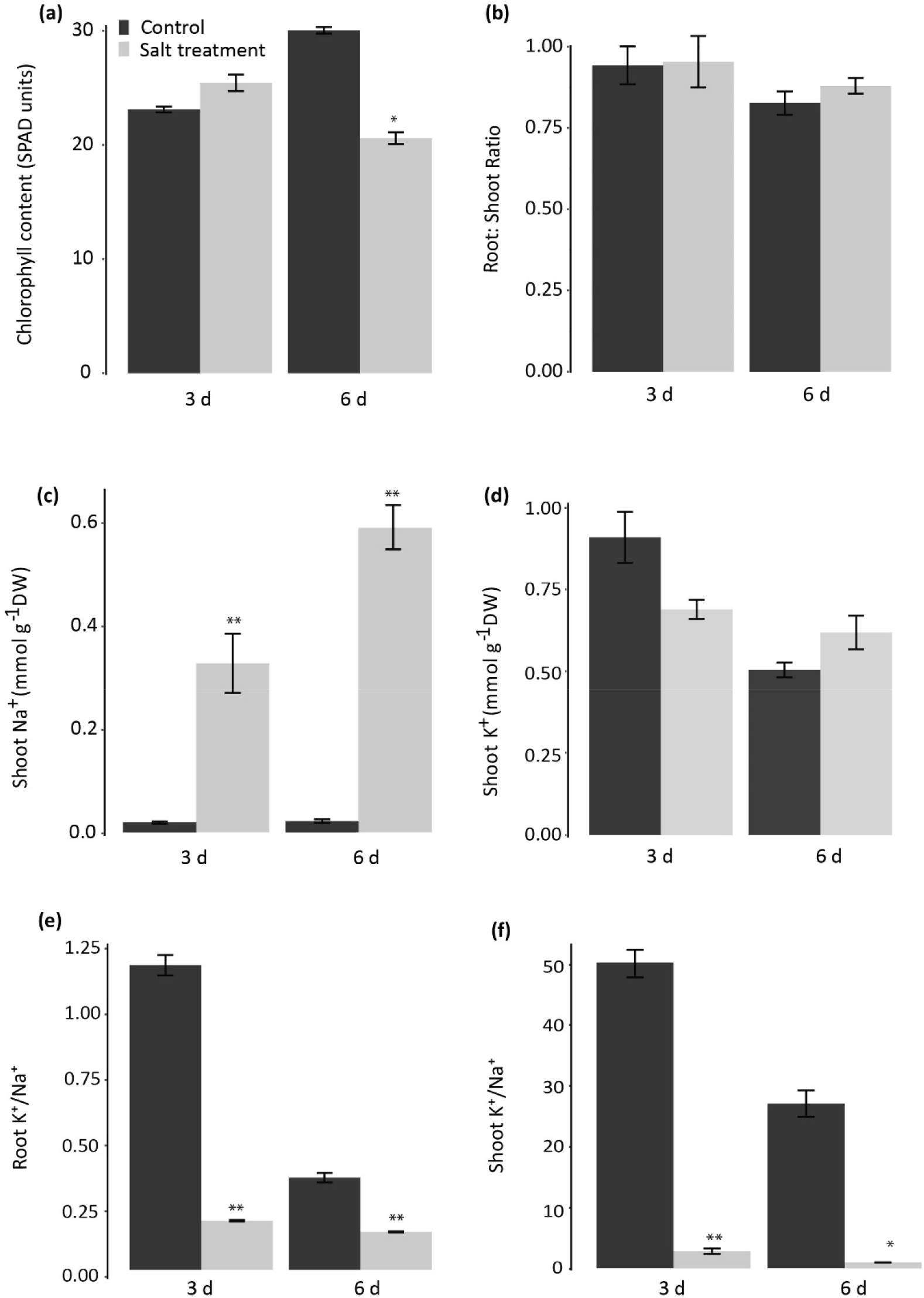
Changes of growth parameters and ion contents in root and shoot tissues of bread wheat cv. Scepter under 150mM NaCl treatment for 3 and 6 days. (a) Chlorophyll content of the fully expanded third leaf, (b) Root:shoot ratio, (c) Na^+^ ion content in shoots, (d) K^+^ ion content in shoots, (e) K^+^/Na^+^ roots, (f) K^+^/Na^+^ shoots. Asterisks indicate the significant difference between control and salt treatment at each single time point separately (Tukey’s Honest Significant Difference; n=4, *, P <0.05)

**Figure S2.**
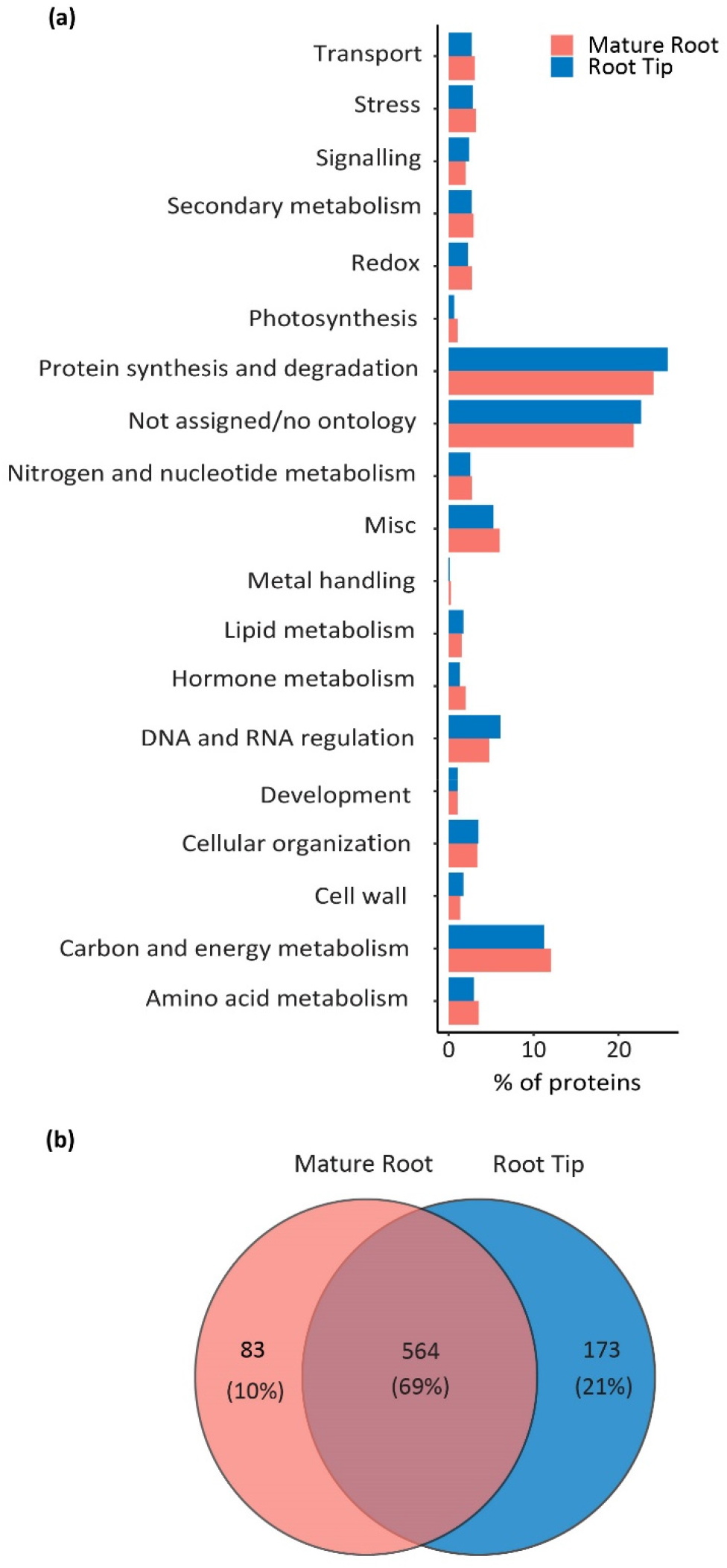
Classification and comparison of the total set of proteins detected by un-targeted proteomics in different bread wheat cv. Scepter root tissues under salt stress. (a) Biological classification of proteins in mature roots and root tips into functional bins defined by Wheat Proteome database, (b) Venn diagram showing the overlap between the total protein sets observed in mature root and root tip

